# Developmental Downregulation of LIS1 Expression Limits Axonal Extension and Allows Axon Pruning

**DOI:** 10.1101/109132

**Authors:** Kanako Kumamoto, Tokuichi Iguchi, Takuya Uemura, Makoto Sato, Shinji Hirotsune

## Abstract

The robust axonal growth and regenerative capacities of young neurons decrease substantially with age. This developmental downregulation of axonal growth may facilitate axonal pruning and neural circuit formation but limits functional recovery following nerve damage. While external factors influencing axonal growth have been extensively investigated, relatively little is known about the intrinsic molecular changes underlying the age-dependent reduction in regeneration capacity. We report that developmental downregulation of LIS1 is responsible for the decreased axonal extension capacity of mature dorsal root ganglion (DRG) neurons. In contrast, exogenous LIS1 expression or endogenous LIS1 augmentation by calpain inhibition restored axonal extension capacity in mature DRG neurons and facilitated regeneration of the damaged sciatic nerve. The insulator protein CTCF suppressed LIS1 expression in mature DRG neurons, and this reduction resulted in excessive accumulation of phosphoactivated GSK-3β at the axon tip, causing failure of the axonal extension. Conversely, sustained LIS1 expression inhibited developmental axon pruning in the mammillary body. Thus, LIS1 regulation may coordinate the balance between axonal growth and pruning during maturation of neuronal circuits.

**Summary Statement:** Developmental downregulation of LIS1 coordinates the balance between axonalelongation and pruning, which is essential for proper neuronal circuit formation but limits nerve regeneration.

## Introduction

The progressive growth of the vertebrate nervous system during embryonic and early postnatal development results in an overabundance of neural connections, requiring targeted elimination to facilitate functional circuit organization (Luo and O’Leary, 2005). An important regressive event is the “pruning” or selective removal of superfluous synapses, axon branches, and dendrites. Axon elimination can occur at different levels, involving small-scale pruning of axon terminals or larger-scale removal of entire collaterals (Vanderhaeghen and Cheng, 2010). Improper pruning in humans has been implicated in various disease states such as autism and schizophrenia (Rosenthal, 2011; Saugstad, 2011). In both flies and mammals, developmental pruning of entire axon branches occurs within a relatively short period of time (Nakamura and O'Leary, 1989; Watts et al., 2003).

Developmental pruning of larger axon segments resembles the fragmentation and eventual disintegration of the distal axon segment following peripheral nerve transection, an active process known as Wallerian degeneration (Conforti et al., 2014). This degeneration allows proximal axons of the peripheral nervous system to regenerate and reinnervate targets, enabling at least partial functional recovery. In contrast to peripheral axons, the repair capacity of central axons is more limited and declines further with age. This unique characteristic of the peripheral nervous system stems at least in part from the ability of Schwann cells (SCs) to provide a proregenerative microenvironment (Son and Thompson, 1995) involving debris clearance, upregulation of membrane-bound and diffusible cues for axonal guidance and organization (Parrinello et al., 2010), and release of prosurvival factors. With advancing age, however, both the speed and the extent of functional recovery decrease following peripheral nerve injury. A number of explanations have been proposed for slower and less complete functional recovery of peripheral axons with age. Adult peripheral axons may regenerate less vigorously than embryonic axons due to intrinsic changes such as decreased axonal transport by cytoskeletal proteins (Brunetti et al., 1987; McQuarrie and Lasek, 1989; Tashiro and Komiya, 1994). Alternatively, older axons may have fewer trophic factor receptors (Ferguson and Son, 2011; Uchida and Tomonaga, 1987). In addition, the periaxonal milieu provided by SCs in older animals has lower concentrations of growth and guidance factors (Bosse, 2012; Komiyama and Suzuki, 1992; Martini, 1994). Finally, impaired neuron-Schwann cell signaling may limit functional recovery by reducing the accuracy of target reinnervation (Kawabuchi et al., 2011). This downregulation of axon regeneration with age may be necessitated by the axon pruning process. One hypothesis is that, to ensure efficient axon pruning, the capacity for axon regeneration must be suppressed.

LIS1 was originally identified as a gene mutated in lissencephaly (Dobyns, 1989; Dobyns et al., 1993; Reiner et al., 1993), a developmental brain disorder caused by defective neuronal migration and consequent cortical dysplasia. Regulation of the motor protein dynein by LIS1 has been intensively investigated (Kardon and Vale, 2009; Vallee and Tsai, 2006; Wynshaw-Boris, 2007). Movement of dynein toward the minus end of the microtubule network is essential for retrograde transport. We previously reported that LIS1 suppresses dynein motility on microtubules in an idling state, which is essential for plus-end-directed (anterograde) transport of dynein by kinesin-1 (Yamada et al., 2008).

Some isolated lissencephaly sequence cases (40%) are caused by *LIS1* haploinsufficiency. We found that *in utero* administration of calpain inhibitors rescued phenotypes of *Lis1*^*+/−*^ mice, including excessive neuronal apoptosis and migration deficits resulting in cortical dysplasia (Yamada et al., 2009). In addition, the blood-brain-barrier- (BBB-) permeable calpain inhibitor SNJ1945, delivered perinatally or *in utero* via pregnant dams, rescued defective neuronal migration in *Lis1*^*+/−*^ mice (Toba et al., 2013).

Here, we demonstrate that physiological LIS1 downregulation via the DNA-binding zinc finger protein CCCTC-binding factor (CTCF) (Ong and Corces, 2014) is responsible for the age-dependent decrease in axonal regenerative capacity of dorsal root ganglion(DRG) neurons. Our findings uncovered a surprising mechanism for the coordination between axonal extension and pruning by the regulation of LIS1 expression.

## Results

### Maturation-Dependent Downregulation of Axonal Extension and LIS1 Expression in DRG Neurons

DRG neurons are a favored model to study axonal regeneration because they possess two axonal branches, one of which projects into the peripheral and the other into the central nervous system. We isolated DRG neurons at various postnatal stages from *Lis1*^+/+^ (wild-type, WT) and *Lis1*^+/−^ (*Lis1*^−/−^ nulls die immediately after implantation) mice (Hirotsune et al., 1998) and examined the potential for axonal extension in cell culture. Axon length of each DRG neuron was defined as the sum of all projections including branches. At postnatal day P3, WT DRG neurons robustly extended axonal processes with extensive arborization, whereas axonal extension was significantly reduced at P15 (Figure 1A, Figure S1A). DRG neurons derived from P3 *Lis1*^+/−^ mice showed reduced axonal extension compared to age-matched WT mice (Figure 1A, Figure S1A), suggesting that LIS1 insufficiency results in earlier downregulation of axonal extension and that LIS1 normally serves to maintain the robust axonal extension capacity of young neurons. Further, we examined whether developmental changes in LIS1 protein expression regulate the neurite extension capacity of DRG neurons. Indeed, LIS1 was induced after plating and the degree of upregulation was age-dependent, with robust induction in P3 WT DRG neurons but much weaker induction at P15 as measured by western blotting (Figure 1B).

**Figure 1:**
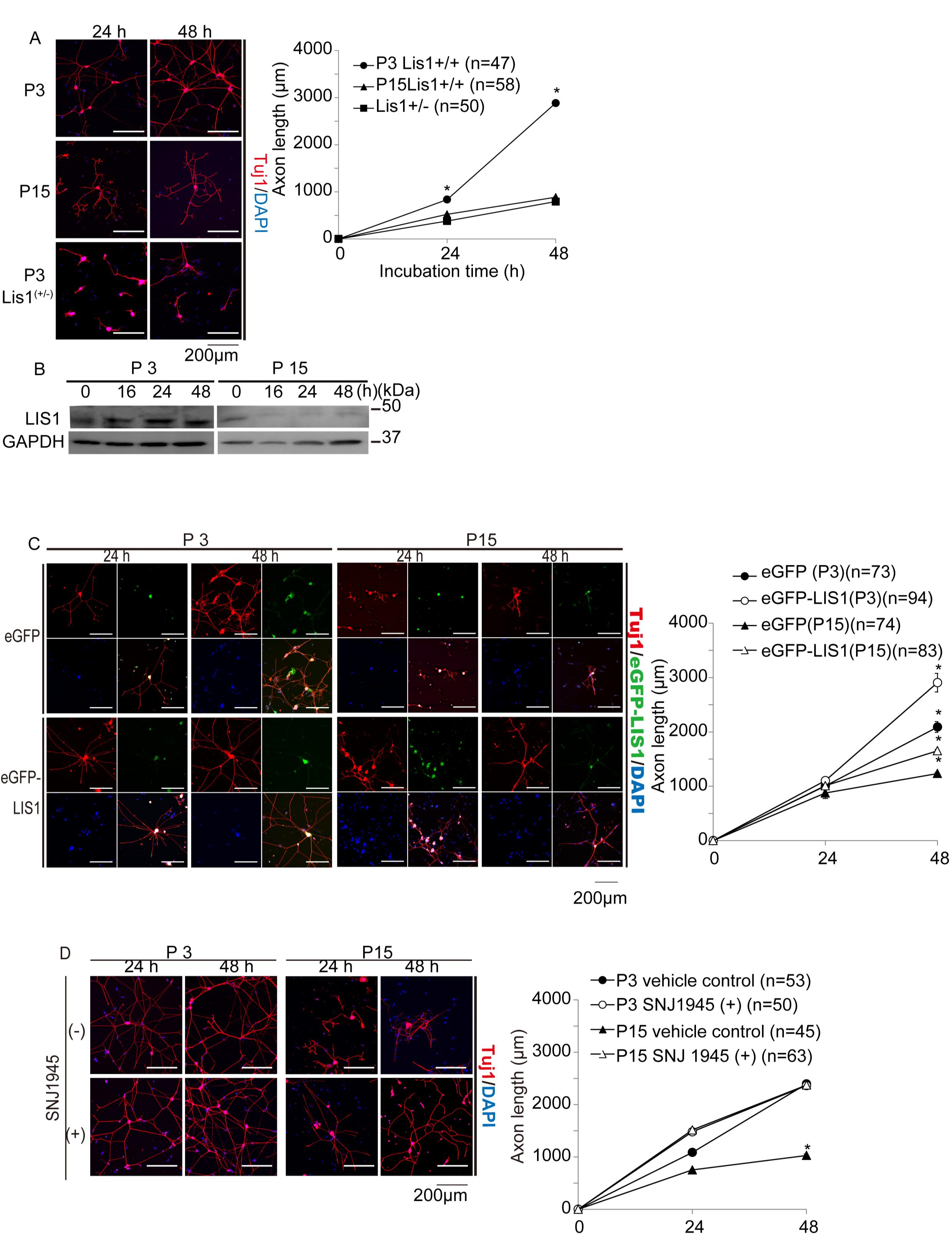
Age-dependent reduction of axonal extension in DRG neurons. Age-dependent downregulation of axonal extension capacity in DRG neurons. Cultures of DRG neurons were isolated from postnatal day P3 and P15 *Lis1*^+/+^ (wild-type, WT) and *Lis1*^+/−^ mice and visualized by Tuj1 immunostaining (red) 24 and 48 h after plating. Nuclei were counterstained with DAPI (blue). Left panels: representative images. Axon length was defined as the summation of all axonal projections including branches. Right panel: average axonal length for each genotype and age with time after plating. Symbols indicate mean axonal lengths with standard errors (mean ± SE). DRG neurons from *Lis1*^+/+^ mice show an age-dependent reduction in axonal extension capacity, while P3 DRG neurons from *Lis1*^+/−^ mice exhibit limited axonal extension capacity of older (P15) *Lis1*^+/+^ neurons. Numbers of neurons examined are indicated in brackets. \**P*< 0.05 by analysis of variance (ANOVA). Left panel: age-dependent LIS1 downregulation in cultured *Lis1*^+/+^ DRG neurons as revealed by western blotting. GAPDH was used as the internal control. Right panel: relative intensities from densitometric analysis. The zero-time LIS1/GAPDH ratio of P3 *Lis1*^+/+^ neurons is defined as 1.0. LIS1 expression is much lower in *Lis1*^+/+^ P15 DRG neurons. ∗*P*< 0.05 by ANOVA. (C) Effect of exogenous LIS1 expression on axonal extension. DRG neurons from P3 and P15 *Lis1*^+/+^ (WT) mice were transfected with eGFP-*Lis1* or empty vector (eGFP). Left panels: representative images. Right panel: quantitation. LIS1 overexpression enhanced axonal extension of DRG neurons at both P3 and P15 compared to age-matched controls (empty vector group). ∗*P*< 0.05 by ANOVA. Effect of the calpain inhibitor SNJ1945 on axonal extension. Left panels: representative images. Right panel: quantitation. SNJ1945 enhanced axonal extension of *Lis1*^+/+^ P15 DRG neurons compared to age-matched vehicle-treated controls but had no effect at P3. ∗*P*< 0.05 by ANOVA.

To confirm that axon extension depends on LIS1 induction, we increased total expression by transfection of enhanced-green-fluorescent-protein- (eGFP-) tagged *LIS1* in WT DRG neurons. Expression of exogenous LIS1 significantly enhanced extension at both P3 and P15 (Figure 1C, Figures S1B and S1C). Under physiological conditions, half of the total LIS1 protein is degraded by calpain-dependent proteolysis at the plus ends of microtubules, and inhibition or knockdown of calpains protected LIS1 from proteolysis and rescued the phenotypes of *Lis1*^+/−^ mice (Yamada et al., 2009). Similarly, peri- or postnatal treatment with the novel calpain inhibitor SNJ1945 rescued defective cortical neuron migration, motor deficits, aberrant neurite length and branch number, and defective retrograde transport of nerve growth factor in *Lis1*^*+/−*^ mice (Toba et al., 2013). Thus, we examined whether endogenous LIS1 augmentation using SNJ1945 (Figure S1E) also facilitates axonal extension. Consistent with exogenous LIS1 overexpression, SNJ1945 significantly enhanced axon extension at P15 (Figure 1D, Figure S1D), although not at P3, possibly due to limited augmentation relative to exogenous overexpression. We conclude that developmental downregulation of LIS1 expression reduces the regenerative capacity of maturing DRG neurons.

### Characterization of the Regulatory Region of *Lis1*

Deletion of the first coding exon of the mouse *Lis1* gene results in the expression of a truncated protein, sLIS1, because of translation initiation at the second methionine (Cahana et al., 2001). Expression of sLIS1 suggests that other regulatory regions may be present outside the vicinity of the first exon, such as within the long first intron. We investigated the transcriptional regulatory region of *Lis1* intron 1 using a *Lis1* minigene conjugated to luciferase as an expression reporter (Figure 2A). Luciferase reporter gene constructs carrying various deletions of intron 1 were transfected into P3 and P15 DRG neurons, followed by the dual-luciferase reporter assay. Deletion construct #59 exhibited higher luciferase activity than the full-length *Lis1* minigene and all deletion constructs, suggesting that a *cis*-repressive element may be present in the deleted region. To narrow down the regulatory region, we created luciferase reporter gene constructs carrying various deletions within the deleted region of the construct #59. The highest luciferase activity was detected in construct #9-1 (Figure 2B), defining the repressor region within a span of 4.5–10 kbp from the start of the first intron. Further, we applied *in silico* prediction of repressors that specifically bind to this region and found a potential binding site for CCCTC-binding factor (CTCF). Therefore, we examined the function of CTCF in the age-dependent repression of LIS1.

**Figure 2:**
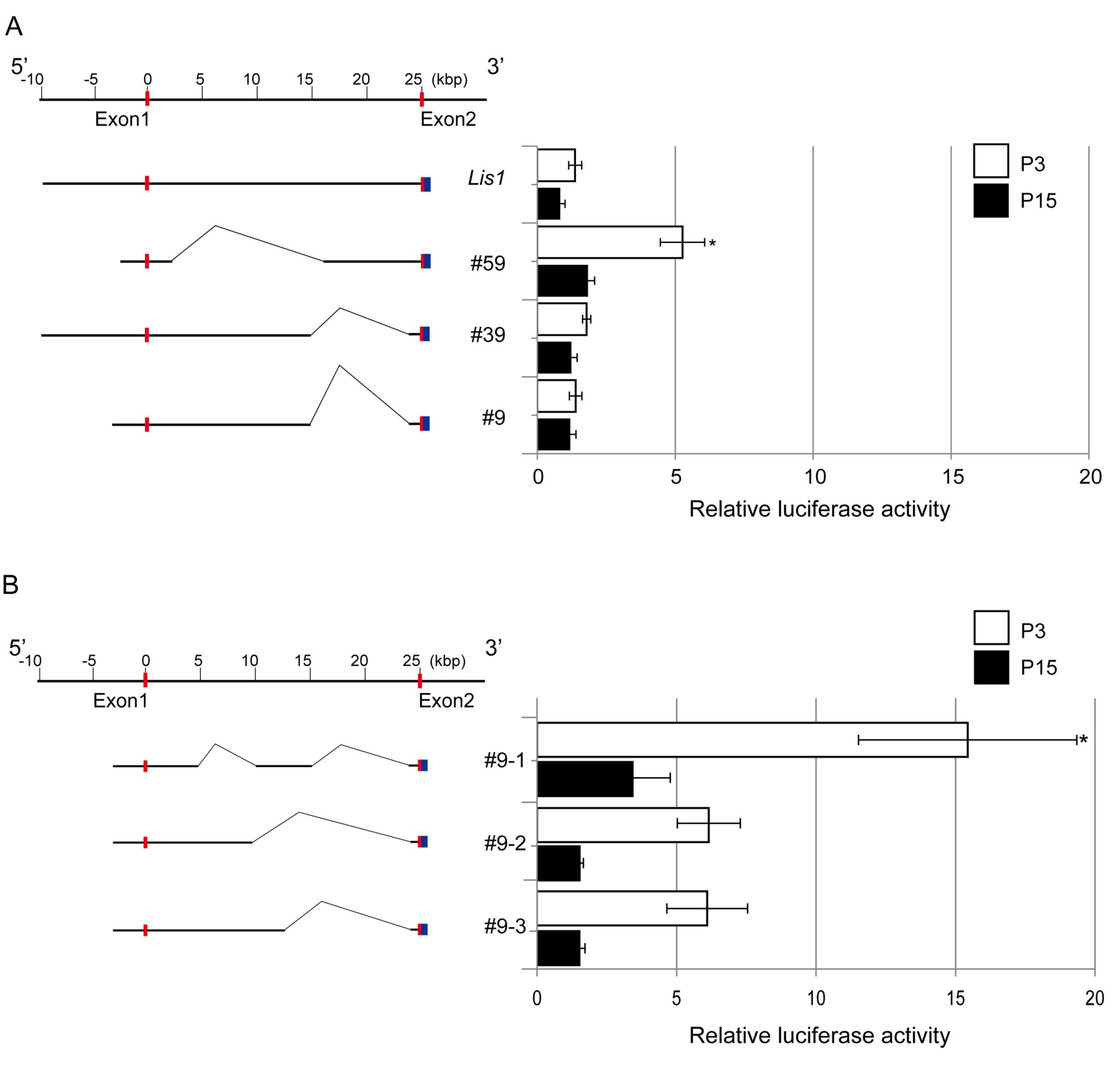
Identification of the *Lis1* suppressor binding site by reporter gene assay. Left panel: Schematic of luciferase reporter constructs containing various deletions within the first intron of *Lis1.* Exons are indicated by red boxes. The luciferase reporter gene was conjugated with *Lis1* exon 2 (blue boxes) in-frame. Constructs were transfected into DRG neurons using the *pGV3* reporter gene vector. The *Renilla* luciferase expression vector *pRLSV40* used as an internal control. Right panel: luciferase activity after 24 h. Relative luciferase activities are shown as mean ± SE of three independent transfected cultures with two replicates per culture. Deletion construct #59 showed the highest luciferase activity, defining the suppressor region to within the deleted span. ∗*P*< 0.05 by ANOVA. (B) Left panel: schematics of constructs including smaller deletions within this span. Right panel: quantitation. Relative luciferase activity was the highest for a deletion of ~4−10 kbp from the start of exon 1 (construct #9-1).

### Negative Regulation of LIS1 Expression by CTCF

CTCF binds to multiple DNA sequences through various combinations of 11 zinc fingers and mediates transcriptional activation/repression and chromatin insulation depending on the biological context (Ong and Corces, 2014). To address whether CTCF suppresses LIS1 expression, we transfected P3 and P15 WT DRG neurons with eGFP-*CTCF* or siRNA against CTCF. Expression of eGFP-CTCF significantly suppressed endogenous LIS1 expression (Figure 3A), whereas CTCF depletion by siRNA enhanced endogenous LIS1 expression (Figure 3B). Therefore, we conclude that CTCF negatively regulates LIS1 expression. Further, exogenous expression of eGFP-CTCF significantly reduced axonal extension at P3 and more mildly suppressed axonal extension at P15 (Figure 3C, Figures S2A and S2B), again consistent with the notion that LIS1 confers greater axonal extension capacity. Moreover, cotransfection of td-Tomato-*Lis1* with eGFP-*CTCF* rescued the decreased axonal extension at P3 (Figure 3C, Figure S2A). On the other hand, CTCF depletion by siRNA transfection in P3 DRG neurons had no effect on axonal extension, whereas depletion at P15 significantly enhanced extension (Figure 4, Figure S3). Thus, we conclude that CTCF controls the age-dependent downregulation of LIS1 and concomitant loss of axonal extension capacity.

**Figure 3:**
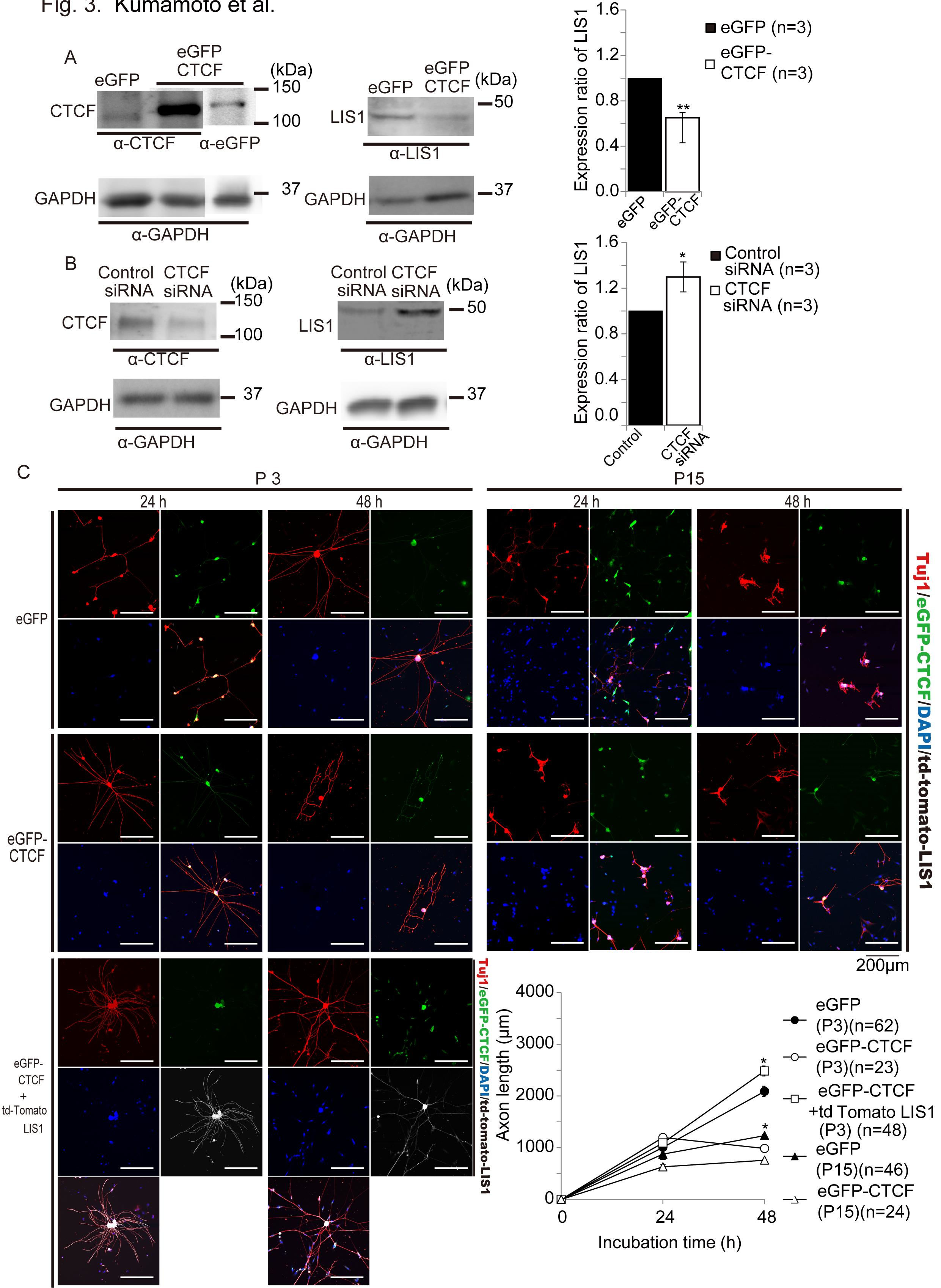
Downregulation of LIS1 expression and axonal extension in WT DRGneurons by CTCF. Effect of exogenous CTCF expression on LIS1 expression in WT DRG neurons was examined by western blotting 48 h after transfection. CTCF overexpression downregulates LIS1. (B) Effect of siRNA-mediated CTCF depletion on LIS1 was examined by western blotting 48 h after transfection. Depletion of CTCF enhances LIS1 expression. GAPDH was used as the internal control. Note: CTCF migrates aberrantly on SDS-PAGE. Endogenous CTCF migrates as a 130 kDa (CTCF-130) protein; however, the open reading frame of the CTCF cDNA encodes only an 82 kDa protein (CTCF-82), in which the N- and C-terminal domains participate in this anomaly (Klenova et al., 1997). Expression of eGFP-*CTCF* was confirmed by western blotting using an anti-GFP antibody (eGFP-CTCF migrated to the same size band as endogenous CTCF protein). Statistical summary of densitometry shown in the right graph. ∗*P*< 0.05 and ∗∗*P*< 0.01 by Student’s *t*-test. (C) Exogenous expression of eGFP-CTCF suppressed axonal extension in WT P3 DRG neurons. Left panels: representative images. Right panel: statistical summary. Suppressive effect of exogenous eGFP-CTCF on axonal extension was rescued by cotransfection with td-Tomato-*Lis1*. ∗*P*< 0.05 by ANOVA.

**Figure 4:**
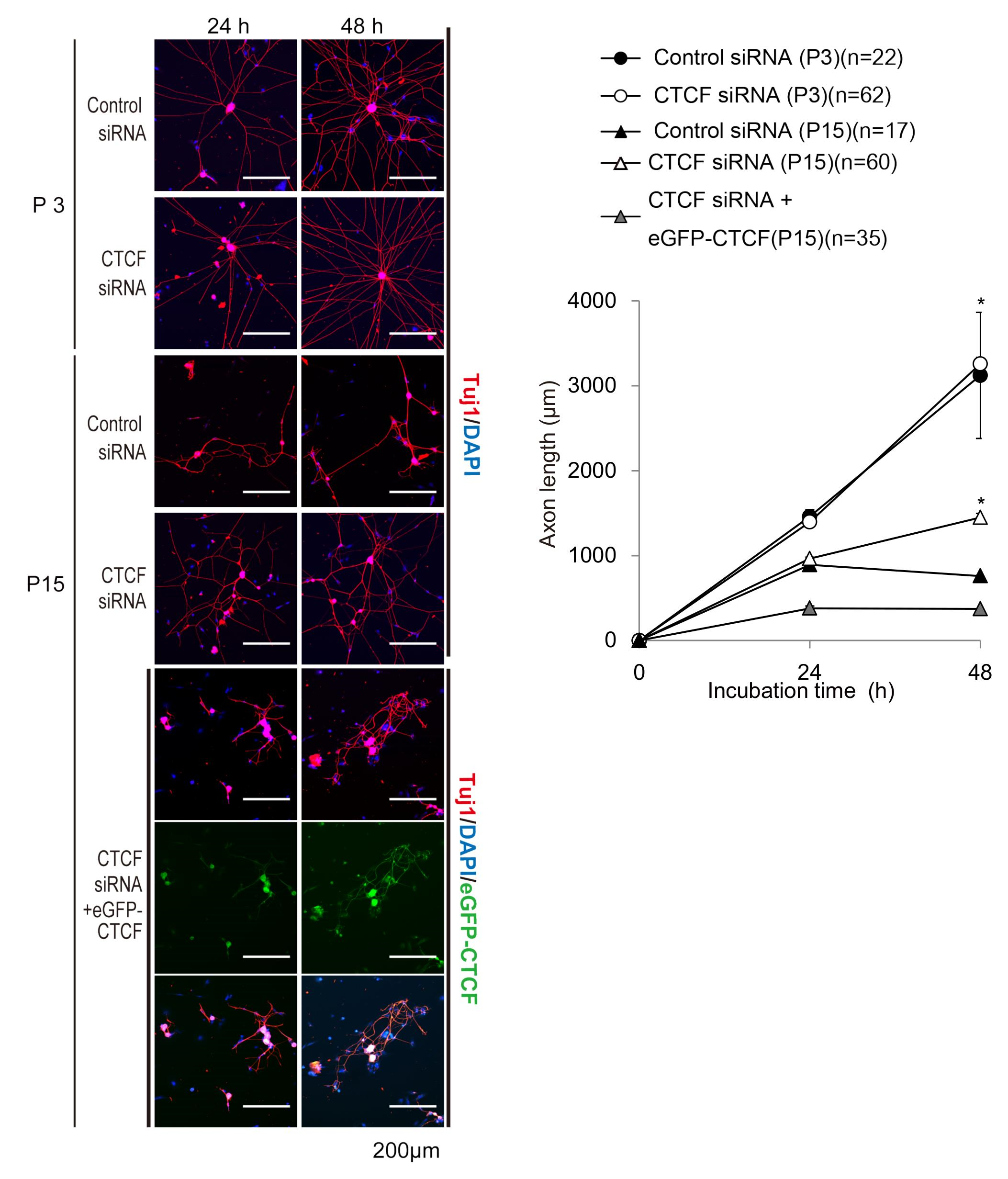
CTCF depletion by siRNA enhances axonal extension capacity at P15. Depletion of CTCF by siRNA enhanced axonal extension in WT P15 DRG neurons, whereas there was no obvious effect on axonal extension in WT P3 DRG neurons (upper panels). Facilitated axonal extension in CTCF-depleted DRG neurons was reversed by cotransfection with eGFP-*CTCF* (lower panels). Representative images shown in left panels, means with standard errors in the right graph. ∗*P*< 0.05 (ANOVA).

### LIS1-Dependent Regulation of GSK-3β Distribution and Activation

GSK3 is a key regulator of neurogenesis, polarization, neurite outgrowth, and plasticity (Hur and Zhou, 2010). GSK3 regulates microtubule growth and stability by phosphorylating microtubule associated proteins (MAPs) such as Tau, MAP1b, CRMP-2, and APC. Mutation or absence of these proteins alters the formation and growth of axons. GSK3-mediated phosphorylation of MAPs such as MAP1B and Tau appears to reduce microtubule binding, thereby creating a population of dynamically unstable microtubules (Trivedi et al., 2005). We previously demonstrated that LIS1 arrests dynein motility and that kinesin-1 transports a LIS1-dynein-tubulin complex to the plus ends of microtubules via mNUDC (Yamada et al., 2010; Yamada et al., 2008). Therefore, we explored whether LIS1 expression modulates GSK3 function via regulation of its dynein-dependent transport. Indeed, GSK-3β was elevated in the growth cones of DRG neurons from *Lis1*^*+/−*^ mice compared to *Lis1*^*+/+*^ mice (Figure 5A), and this aberrant accumulation of GSK-3β was reversed by administration of SNJ1945 (Figure 5A). These results suggest that elevated LIS1 expression in young neurons serves to prevent GSK-3β accumulation in growth cones, concomitant MAP phosphorylation, and ensuing destabilization of microtubule dynamics, thereby allowing for axonal extension.

**Figure 5:**
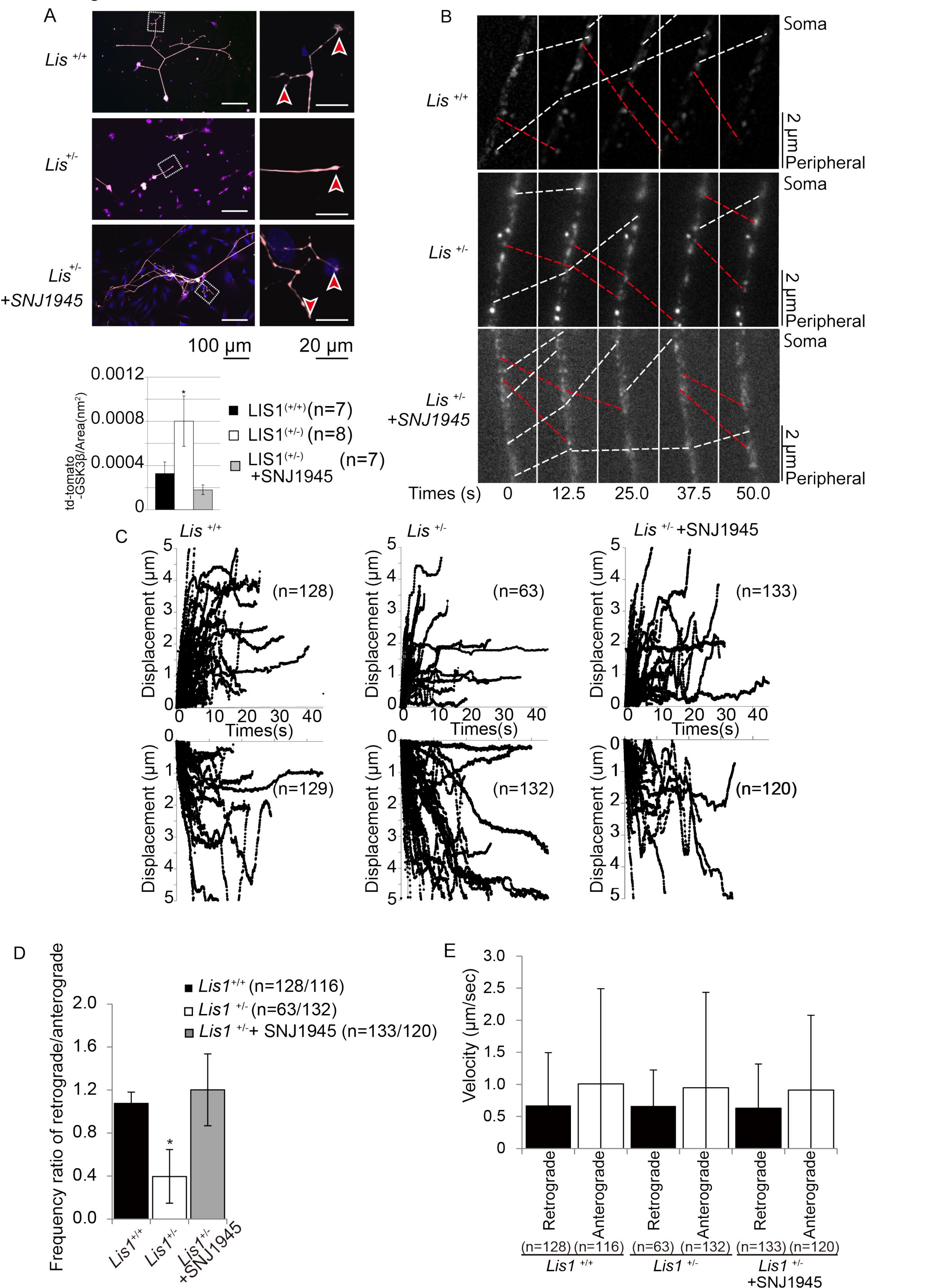
Axonal transport of GSK-3β in DRG neurons. Distribution of GSK-3β in P3 DRG neurons from *Lis1*^*+/+*^ (WT), untreated *Lis1*^*+/−*^, and SNJ1945-treated *Lis1*^*+/−*^ mice. Red arrowheads indicate growth cones of DRG neurons. Quantitation of fluorescence intensity (bottom) reveals GSK-3β accumulation in growth cones of *Lis1*^*+/−*^ mice compared to *Lis1*^*+/+*^ mice and rescue of aberrant growth cone accumulation by SNJ1945. ∗*P*< 0.05 (ANOVA). (B) eGFP-*GSK-3*β was expressed in P3 DRG neurons and monitored by time-lapse fluorescence microscopy. Anterograde movement and retrograde movement are shown by red dotted lines and white dotted lines, respectively. Elapsed time is indicated at the bottom. (C) Trajectories of eGFP-GSK-3β movement in axons of P3 DRG neurons from *Lis1*^*+/+*^ mice (left panels), *Lis1*^*+/−*^ mice (middle panels), and SNJ1945-treated *Lis1*^*+/−*^ mice (right panels). Retrograde and anterograde movements are shown in upper and lower panels, respectively. Note the diminished retrograde displacement (upper middle panel) in untreated *Lis1*^*+/−*^ mice. (D) The ratio of retrograde frequency to anterograde frequency of eGFP-GSK-3β in DRG neurons. Retrograde movement frequency was significantly lower in *Lis1*^*+/−*^ mice and rescued by SNJ1945. ∗*P*< 0.05 by ANOVA. (E) Velocity of eGFP-GSK-3β axonal transport in P3 DRG neurons. There was no significant difference in transport velocity during displacement between *Lis1*^*+/−*^ and *Lis1*^*+/+*^ mice.

To determine whether this GSK-3β accumulation occurs via LIS1 effects on axonal transport, we conducted live cell imaging of P3 DRG neurons expressing td-Tomato-tagged GSK-3β. The fusion protein exhibited robust bidirectional movement, suggesting that the subcellular distribution of GSK-3β relies on the activity of motor proteins, including kinesin and dynein. While there was no significant difference in the speed of retrograde GSK-3βmovement between *Lis1*^*+/−*^ and *Lis1*^*+/+*^ axons (Figures 5B, 5C, and 5E, Movie S1), the ratio of retrograde to anterograde movement was significantly lower in DRG neurons from *Lis1*^*+/−*^ mice (Figures 5B, 5C, and 5D, Movie S1). This decreased frequency of retrograde movement in *Lis1*^*+/−*^ DRG neurons was rescued by the addition of SNJ1945 (Figures 5B–5D, Movie S1). We conclude that LIS1 expression is essential for the retrograde transport of GSK-3β and prevention of distal accumulation in growth cones.

Phosphorylation of GSK-3βat Ser9 renders it inactive (Dudek et al., 1997), whereas phosphorylation at Tyr216, which lies within the activation loop between subdomains VII and VIII of the catalytic domain, is necessary for functional activity (Hughes et al.,1993). A constitutively active GSK-3β mutant inhibited axon formation, whereas multiple axons formed from a single neuron when GSK-3β activity was reduced by small molecule inhibitors, a peptide inhibitor, or siRNAs (Jiang et al., 2005). To estimate the level of GSK-3β activity at the tip of the DRG neurons, we double-labeled DRG neurons from *Lis1*^*+/−*^ and *Lis1*^*+/+*^ mice with an antibody against total GSK-3β and another against either (inactive) GSK-3β phosphorylated at Ser9 (pS9-GSK-3β) (Figure 6) or active GSK-3β phosphorylated at Tyr 216 (pY216-GSK-3β) (Figure 7) and determined the inactive/total and active/total GSK-3β ratios. The pS9-GSK-3β/GSK-3β ratio in the growth cones of P3 DRG neurons from *Lis1*^*+/−*^ mice was significantly lower than in growth cones of P3 *Lis1*^*+/+*^ mice (Figures 6A and 6B, and 6D). Conversely, the pY216-GSK-3β/GSK-3β ratio in the growth cones of P3 DRG neurons from *Lis1*^*+/−*^ mice was significantly higher than in growth cones of P3 *Lis1*^*+/+*^ mice (Figures 7A, 6B, and 6D). Further, this aberrant accumulation of GSK-3β in *Lis1*^*+/−*^ mice was rescued by SNJ1945 (Figures 6C and 6D, Figures 7C and 7D). Collectively, we conclude that GSK-3β is overactivated in the growth cones of P3 DRG neurons from *Lis1*^*+/−*^ mice, which will suppress axonal extension.

**Figure 6:**
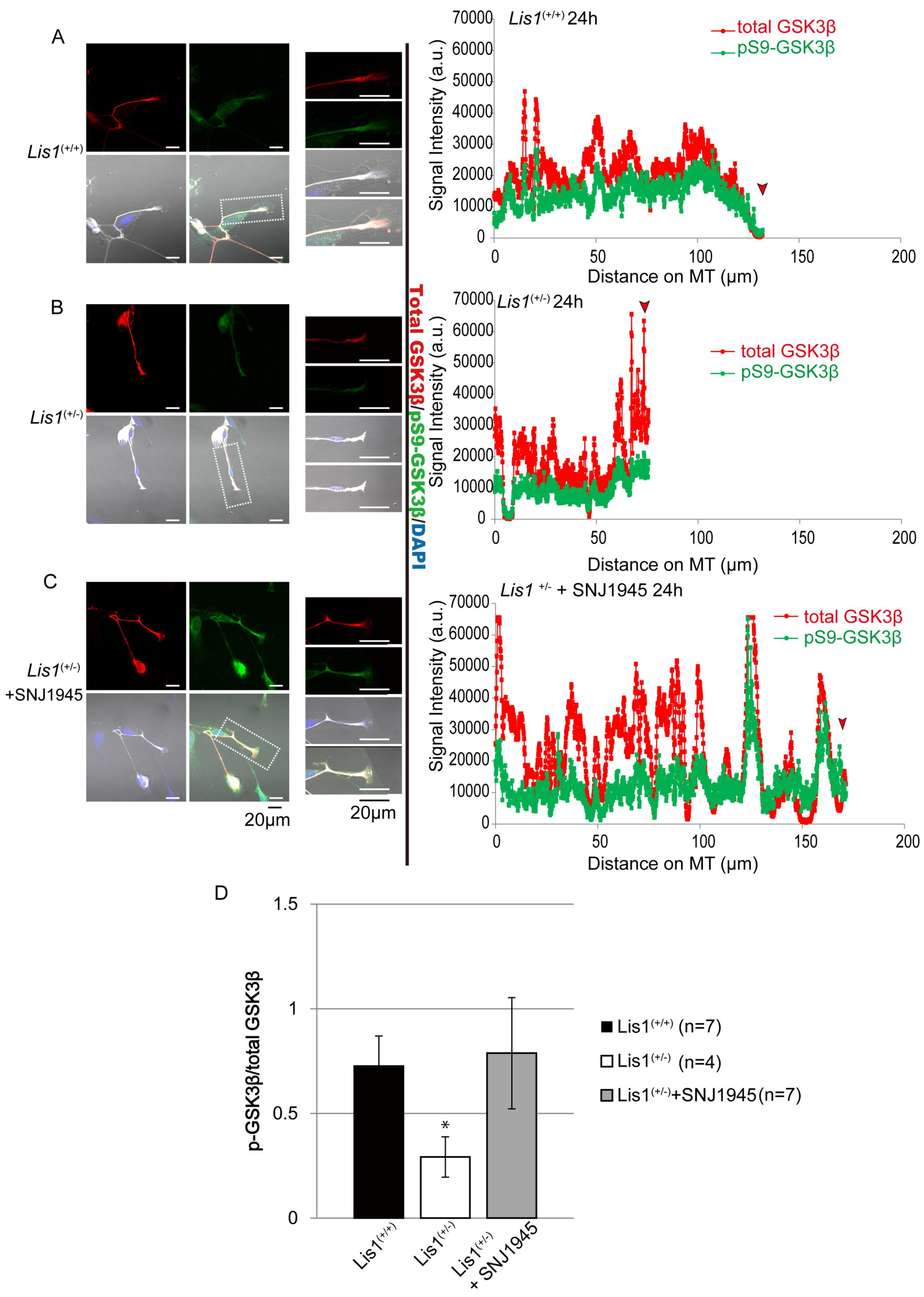
Distribution of phospho-GSK-3β (Ser9) in DRG neurons. (A−C) Left panels: DRG neurons from (A) *Lis1*^*+/+*^ mice, (B) *Lis1*^*+/−*^ mice, and (C) SNJ1945-treated *Lis1*^*+/−*^ mice stained with anti-GSK-3β (red) and anti-pS9-GSK-3β (green). Middle panels: higher-magnification images of an area demarcated in the left panels (white boxes). Right panels: normalized fluorescence intensity along the axon. Red arrowheads indicate growth cones. Greater fluorescence intensity at the growth cones of DRG neurons from *Lis1*^*+/+*^ (middle) and SNJ1945-treated *Lis1*^*+/−*^ (bottom) mice indicates accumulation of total GSK-3β and anti-pS9-GSK-3β. (D) Quantitation of anti-pS9-GSK-3β to total GSK-3β intensity in growth cones indicates relatively lower accumulation of the inactive anti-pS9-GSK-3β in *Lis1*^*+/−*^ growth cones and reversal by SNJ1945. ∗*P*< 0.05 (ANOVA).

**Figure 7:**
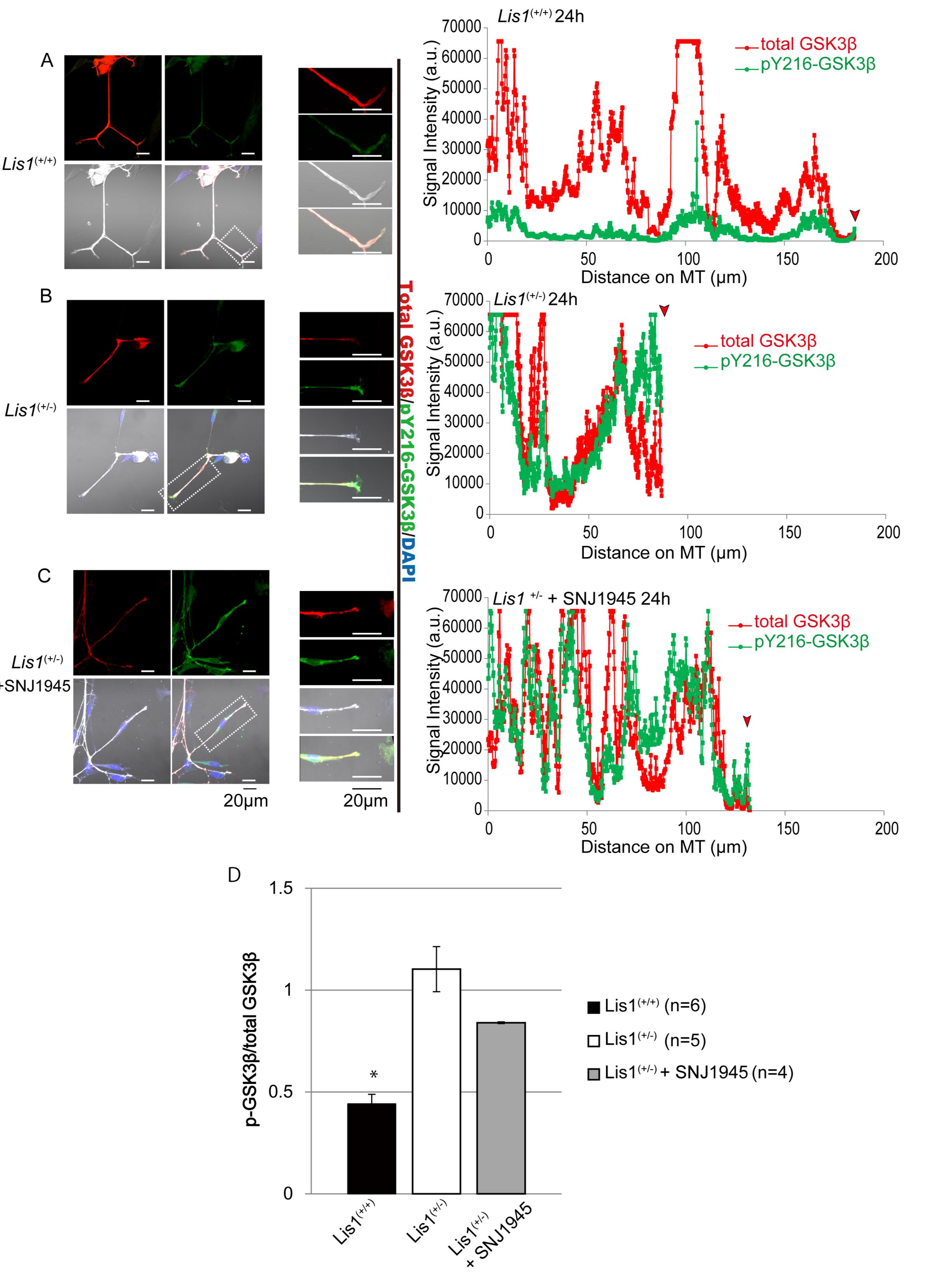
Distribution of phospho-GSK-3β (Tyr216) in DRG neurons. (A−C) Left panels: DRG neurons from (A) *Lis1*^*+/+*^ mice, (B) *Lis1*^*+/−*^ mice, and (C) SNJ1945-treated *Lis1*^*+/−*^ mice. Anti-GSK-3β and anti-pY216-GSK-3β immunostaining. Middle panels: higher-magnification images of an area demarcated in the left panels (white boxes). Left panels: normalized fluorescence intensity along the axon length. The red arrow indicates the position of the growth cone (tip). (D) Quantitation of anti-pS9-GSK-3β to total GSK-3β intensity in growth cones indicates relatively greater accumulation of the active pY216-GSK-3β form in *Lis1*^*+/−*^ growth cones compared to*Lis1*^*+/+*^ growth cones.∗*P* < 0.05 by ANOVA.

### Facilitation of Axonal Regeneration by SNJ1945 after Sciatic Nerve Injury

We previously reported that a calpain inhibitor upregulates LIS1 (Yamada et al., 2009) and facilitates neuronal circuit formation (Toba et al., 2013). To investigate whether endogenous LIS1 augmentation using SNJ1945 facilitates axonal regeneration after injury, we subjected mice to sciatic nerve (SN) axotomy, a common model of peripheral nerve injury in rodents (Magill et al., 2007), and compared regrowth and functional recovery between SNJ1945-treated and untreated groups. The left SN was transected at the obturator tendon level in 4-week-old WT mice, and the extent of injury and subsequent regeneration were evaluated by light and electron microscopy. Following nerve injury, toluidine blue staining of nerve cross sections revealed changes in the distal nerve stump characteristic of Wallerian degeneration, such as massive axonal swelling. Treatment with SNJ1945 had no obvious effect on SN regeneration at one week (Figures 8A and 8B), but treated mice exhibited significantly more numerous myelinated SN fibers at one month (Figure 8C) and numerically greater numbers at three months (Figure 8D) and six months (Figure 8E) after transection (results summarized in Figure 6F). Transected SNs from control mice exhibited only partial regeneration with hypomyelination compared to SNJ1945-treated mice at the same times after injury. This suggests that SNJ1945-induced enhancement of axonal L1S1 accelerates regeneration of the injured sciatic nerve (Figures 8C–8F).

**Figure 8:**
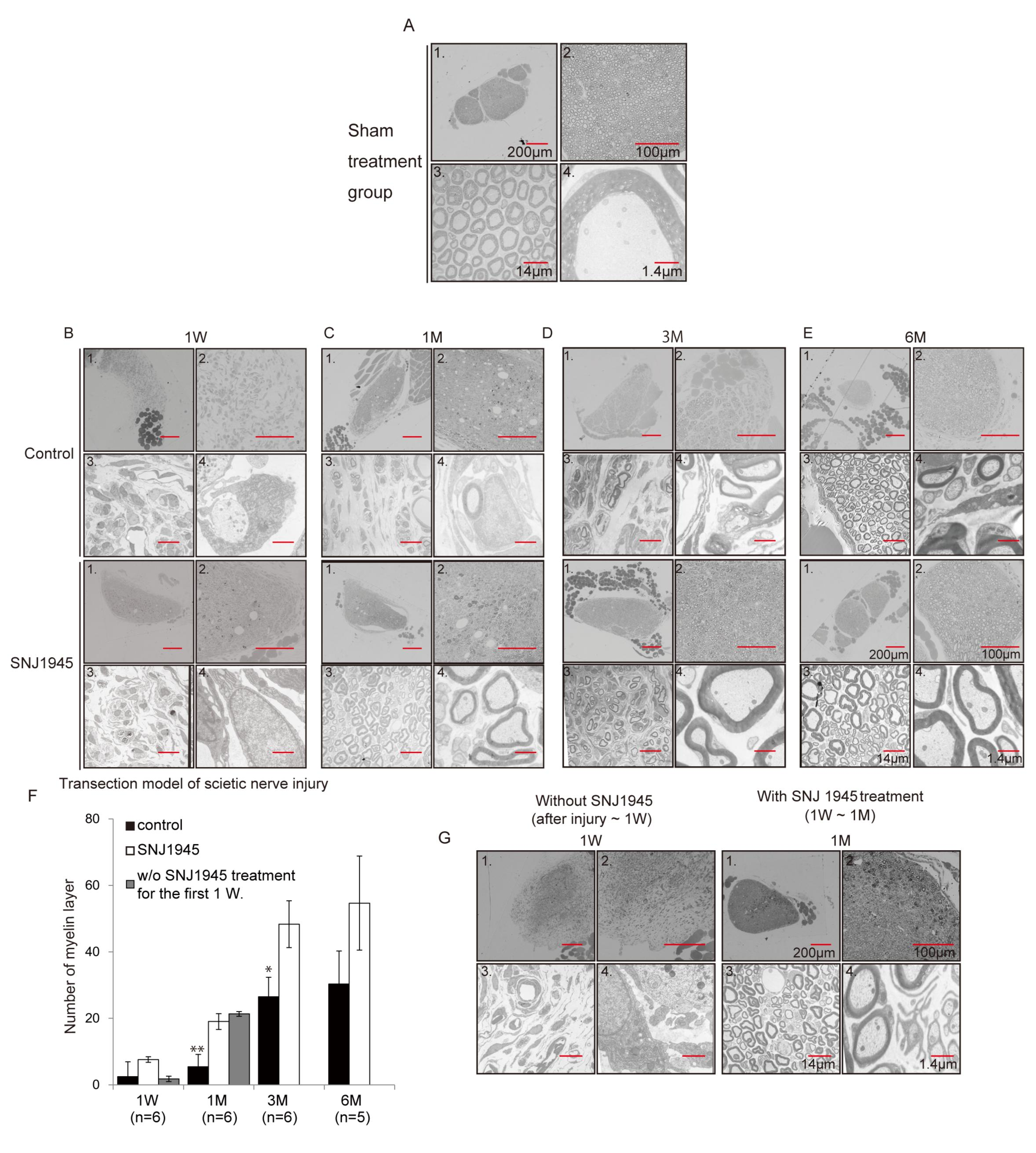
Facilitation of sciatic nerve regeneration by SNJ1945. Transection model of mouse SN injury. (A) Control images of SN cross sections from sham-treated WT mice. Imaging modality, orientation, and magnification are as follows: panel 1, low-magnification Prussian blue staining; panel 2, high-magnification Prussian blue staining; panel 3, electron microscopic (EM) image at low magnification; panel 4, EM at higher-magnification. Effects of oral SNJ1945 treatment on SN axon remyelination one week (B), one month (C), three months (D), and six months (E) after transection. Upper panels are from the untreated transection group (control). Lower panels are from mice treated with oral SNJ1945 after SN transection. SNJ1945 treatment accelerated remyelination of the sciatic nerve. (F) Statistical summary (mean ± SE for 10 mice) showing enhanced numbers of regenerated myelinated SN axons in SNJ1945-treated mice after transection. ∗*P*< 0.05 and ∗∗*P*< 0.01 (ANOVA). (G) Effect of SNJ1945 on regeneration/remyelination when administration was delayed for one week after transection. Left panels: one week after transection (without treatment). Right panels: after one month of treatment. SNJ1945 treatment was still effective.

In principle, this accelerated regeneration after injury could be mediated by facilitation of proximal axon growth, protection against proximal regression, accelerated distal Wallerian degeneration, or a combination. Previous studies examining the sequence of events following injury revealed at least three morphologically discernible phases (Wang et al., 2012). Seventy-two hours after transection, rapid fragmentation and cytoskeletal breakdown occurred along the full length of the distal axon, followed by increased microglial influx to clear axonal remnants. So, to distinguish among these possible mechanisms, we initiated SNJ1945 treatment one week after transection, beyond the early phase of degeneration. Although this latency attenuated initial recovery and remyelination of transected SN at one week, remyelination was still significantly augmented at one month (Figure 8F and 8G). We conclude that SNJ1945 treatment facilitates axon regeneration rather than removal of distal debris.

To examine the efficacy of SNJ1945 treatment on functional motor recovery, walking-track analysis was performed using the sciatic function index (SFI) (de Medinaceli et al., 1984). Control transection model mice displayed prolonged functional deficits (Figures 9A–9C, Movies S2 and S3) and significantly lower SFI values throughout the twelve-week assessment period compared to SNJ1945-treated mice (Figures 9A–9C, Movies S2 and S3). These results suggest that SNJ1945 promotes SN reinnervation of appropriate muscle targets for motor function recovery but upregulating LIS1 expression.

**Figure 9:**
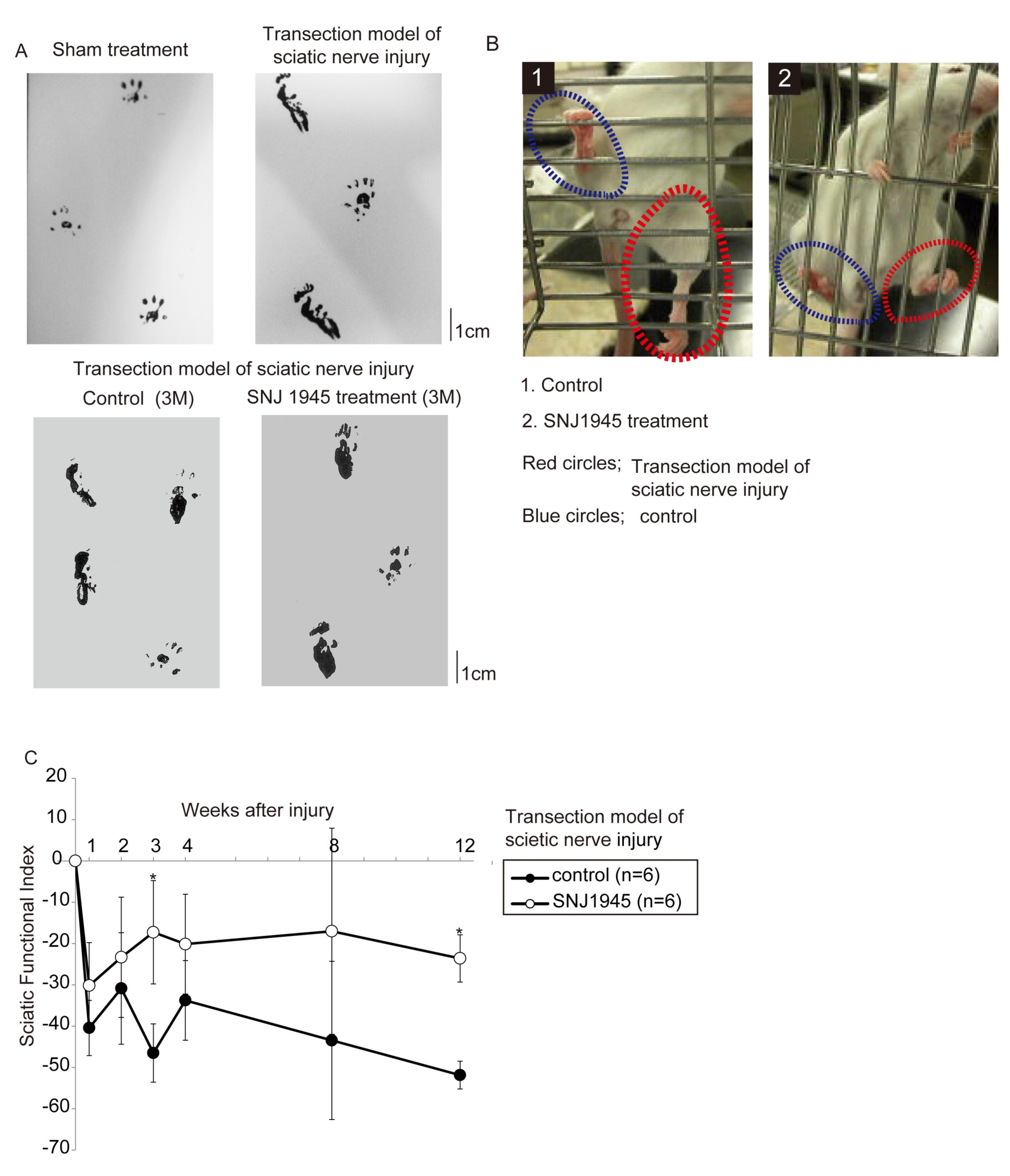
Gait analysis during recovery from SN transection. Footprint patterns of mice following unilateral SN transection. Untreated model mice exhibited imbalanced and asymmetric gait patterns with increased “toe-out” angles in the injured limb and asymmetric right versus left limb step lengths. SNJ1945 treatment improved asymmetric gaiting. (B) Photos of mouse legs after three month of the drug treatment. The control mouse displayed defective grasping of the cage bars by the foot on the injured (left) side, whereas the SNJ1945-treated mouse displayed partial improvement (right side). (C) SFI of gait analysis indicating facilitated functional recovery after transection in SNJ1945-treated mice. ∗*P*< 0.05 by Mann–Whitney *U* test.

### Axonal Pruning and LIS1 Downregulation

Immature neuronal networks formed by axonal and dendritic sprouting subsequently undergo extensive pruning to form functional circuits. This pruning includes distinct processes for removal of axons, axon branches, and dendrites (Luo and O’Leary, 2005). It is speculated that, to ensure efficient axonal pruning, the capacity for axonal regeneration must be suppressed. We hypothesized that developmental LIS1 downregulation may be associated with axon and dendrite pruning because pruning and growth/regeneration require reciprocal effects on growth-associated processes such as cytoskeletal dynamics.

We first examined the association of axonal extension during cortical development with LIS1 expression. Like DRG neurons, cortical neurons isolated from P3 mice exhibited robust axonal outgrowth, whereas cortical neurons isolated from P15 or P60 mice showed markedly reduced outgrowth potential (Figure 10A). Thus, as in peripheral neurons, LIS1 downregulation was coupled to the maturation of cortical neurons (Figure 10B). We conclude that the parallel age-dependent reduction of axonal growth potential and LIS1 expression observed in the DRG is recapitulated in the central nervous system.

**Figure 10.**
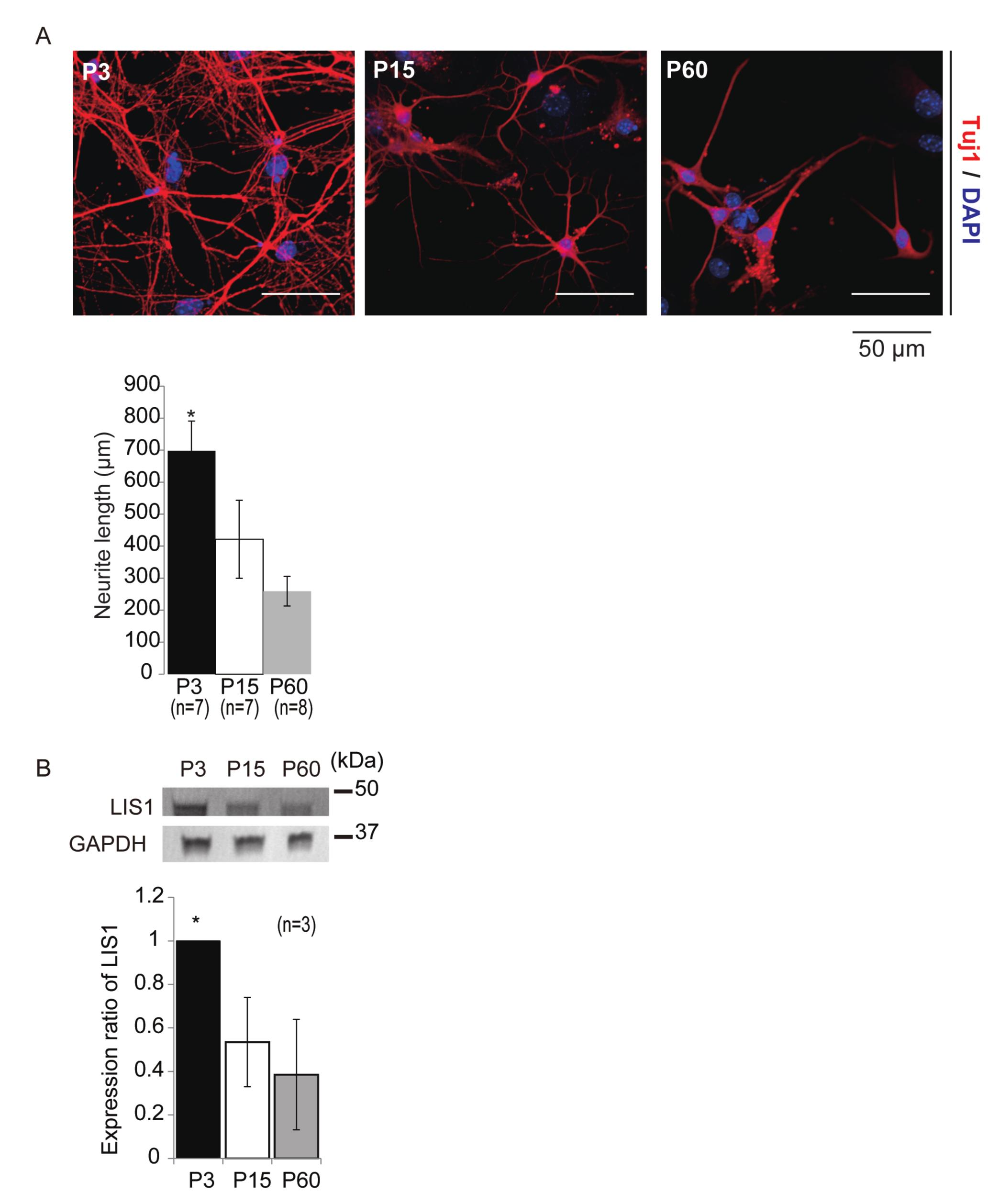
Regulatory function of LIS1 and CTCF during embryonic brain development. (A) Age-dependent downregulation of axonal extension capacity in cortical neurons. Cortical neurons were isolated from WT P3, P15, and P60 mice. Axons were visualized by Tuj1 immunostaining (red) 7 days after plating. Nuclei were counterstained with DAPI (blue). Upper panels: representative images. Lower panel: mean (±SE) axonal extension revealing age-dependent reduction as in DRG neurons. Numbers of neurons examined indicated in brackets. ∗*P*< 0.05 by ANOVA. (B) LIS1 expression in the brain examined by western blotting with GAPDH as the internal control. Endogenous expression of LIS1 was downregulated at P15 and P60.

We next examined neuronal circuit maturation *in vivo* under modulation of LIS1 or CTCF expression to assess effects on pruning. We focused on the postmammillary component of the mouse fornix, a tract of axons that extends beyond the mammillary bodies and into the midbrain in the first postnatal week as the fornix-mammillary projection is established but progressively regresses until it is no longer detectable by the third postnatal week (Stanfield et al., 1987). To visualize the developing fornix, bacterial artificial chromosome (BAC) transgenic mice (Fezf2-Gfp), in which GFP expression is regulated by the promoter for the transcription factor Fezf2 (Gong et al., 2003; Kwan et al., 2008), were transfected with control, LISI, CTCF, or CTCF-targeted shRNA expression vectors by in utero gene transfer (Tabata and Nakajima, 2001). Transfection of a red fluorescent td-Tomato control plasmid at embryonic day 12.5 (E12.5)(Figures 11A and 11B) revealed numerous labeled fornix fibers extending into and beyond the mammillary body at P15. However, this postmammillary population was markedly diminished by P18 (Figure 11B) and completely absent at P21 (Figures 11B and 11F). However, a substantial fraction of the postmammillary component survived at P21 in mice transfected with td-Tomato-*Lis1*, with the labeled axons confined to a sharply defined fiber bundle at the dorsolateral aspect of the mammillary nuclei (Figures11C and 11F). Similarly, when CTCF was depleted by a targeted shRNA, survival of the postmammillary component was significantly increased at P21 compared to controls, with labeled fascicles continuing into the mammillary body (Figures 11D and 11F). (Alant et al., 2013)In contrast, the postmammillary component underwent premature regression in mice overexpressing td-Tomato-*CTCF*, as surviving fascicles extending into the mammillary body were clearly diminished at P15 (Figures 11E and 11F). Thus, CTCF-dependent LIS1 suppression is critical for axonal regression during development.

**Figure 11.**
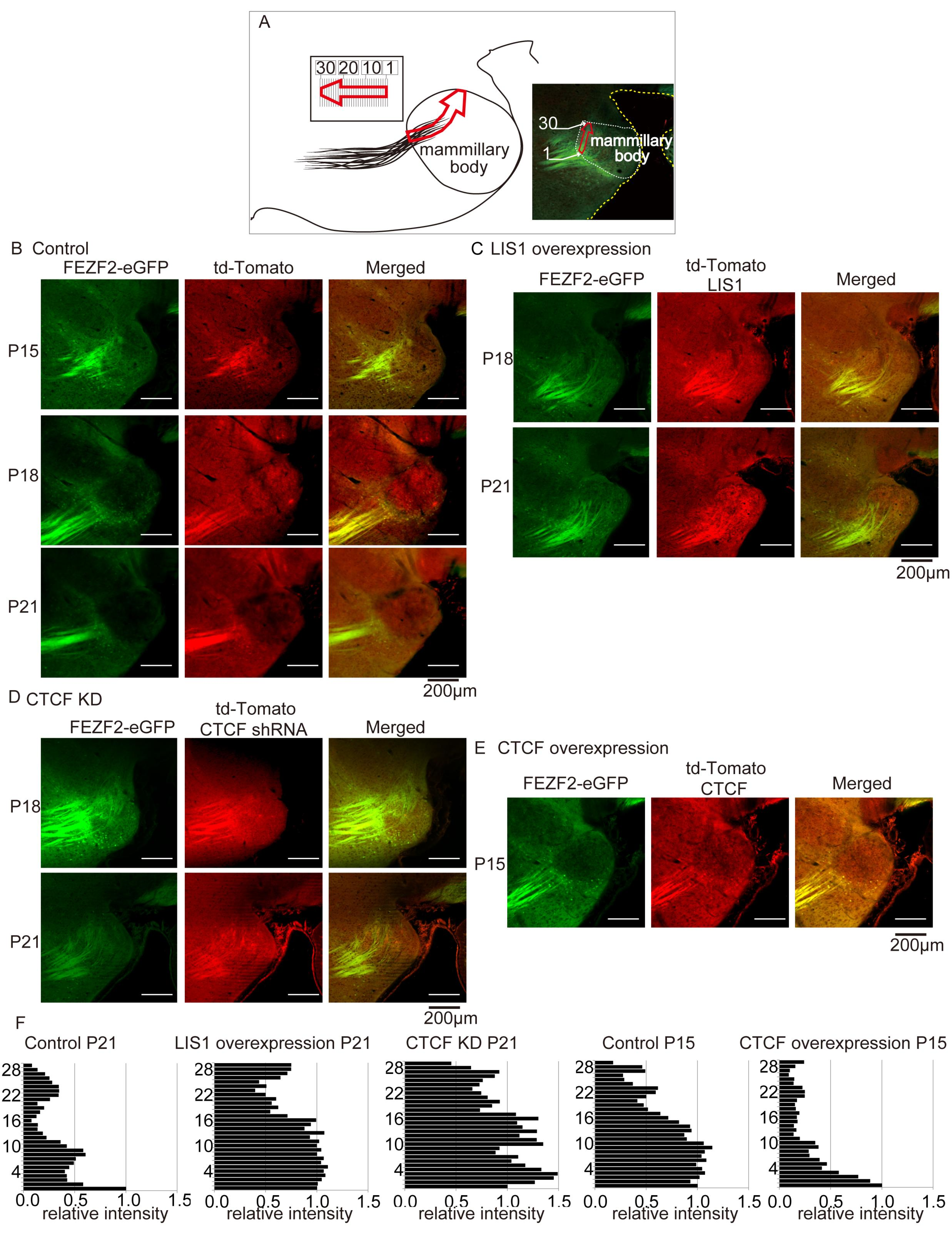
Developmental LIS1 downregulation promotes axonal pruning of the postmammillary component of the fornix. (A) CAG promoter-driven td*-Tomato* was expressed in *Fezf2-Gfp* BAC transgenic mice to trace the postmammillary component of the fornix. Illustration indicates the orientation of the postmammillary component, which was divided into 30 bins for quantitation. (B) CAG-*td-Tomato* expression vector was electroporated at E12.5 and mice were inspected at P15, P18, and P21. The postmammillary component was still present at P15, pruned by P18, and completely absent at P21. (C) CAG-td*-Tomato-Lis1* was introduced at E12.5 and the postmammillary component examined. The postmammillary component was still present at P18 and P21, indicating that LIS1 overexpression suppressed pruning of the postmammillary component. (D) shRNA against *CTCF* was introduced at E12.5 and the postmammillary component examined. The postmammillary component was still present at P18 and P21. (E) CAG-td*-Tomato-CTCF* was introduced at E12.5 and the postmammillary component examined. Postmammillary component disappeared prematurely by P15. (F) Quantitation of fluorescence intensity in each bin of the postmammillary component (see Figure A).

## Discussion

We provided evidence that endogenous LIS1 is a key regulator of the balance between neural circuit formation and regenerative capacity in both the developing peripheral and the central nervous systems. DRG neurons exhibited an age-dependent reduction in axonal extension potential that was paralleled by downregulation of LIS1 expression. Further, extension potential was also reduced in young DRG neurons from *Lis1*^*+/−*^ mice. In both P15 WT and P3 *Lis1*^*+/−*^ neurons, low extension potential was rescued by exogenous LIS1 overexpression and by endogenous LIS1 augmentation using the calpain inhibitor SNJ1945. In addition, we show that LIS1 is physiologically regulated by the insulator protein CTCR. LIS1 augmentation by SNJ1945 (Toba et al., 2013) promoted axonal extension, which suggests a novel approach for improving peripheral nerve regeneration after injury. A recent report provided evidence that the endogenous calpain inhibitor calpastatin functions as a determinant of axonal survival both during development and after injury (Yang et al., 2013). Therefore, calpastatin induction is a promising approach for blocking LIS1 suppression by calpains, thereby reinducing robust axonal growth. Oral SNJ1945 treatment in particular is a promising approach because SNJ1945 has low toxicity and high BBB permeability (Toba et al., 2013). Indeed, oral SNJ1945 markedly accelerated recovery of motor function following sciatic nerve transection in mice.

We also demonstrated that CTFC is involved in the physiological downregulation of LIS1. CTCF contains a highly conserved DNA-binding domain with 11 zinc fingers, enabling it to function as a major insulator for numerous target genes. In fact, CTCF is present at 55,000–65,000 sites in the mammalian genome (Ong and Corces, 2014). Depletion of CTCF at P15 facilitated axonal extension through LIS1 overexpression, whereas CTCF overexpression at P3 suppressed axonal extension by LIS1 downregulation. In contrast, however, individual projection neurons in CTCF-cKO mice exhibited significantly reduced average dendritic lengths (Hirayama et al., 2012) rather than overextension. This contradictory result may indicate distinct regulatory mechanisms for axons and dendrites or arise because of different KO procedures. In the NEX-Cre mice used by Hirayama et al. (2012) for disruption of CTCF in postmitotic neurons, the most prominent Cre activity was observed in the neocortex and hippocampus beginning at around E11.5 (Goebbels et al., 2006). Within the dorsal telencephalon, Cre-mediated recombination was substantial in hippocampal pyramidal neurons, dentate gyrus hilar mossy cells, and DR granule cells, but absent from proliferating neural precursors (Goebbels et al., 2006). In the current study, an shRNA against CTCF was transfected into E12.5 embryos to examine the effect of depletion of CTCF on postmammillary fornix pruning. Thus, the discrepancy may be attributable to the earlier developmental stage of CTCF disruption, which in turn could influence numerous subsequent developmental processes. Alternatively, CFCT functions in neural development may be region-specific.

We found that LIS1 regulates axonal extension via transport of GSK-3β. GSK-3β is amultifunctional serine/threonine kinase known to regulate axon growth (Hur and Zhou, 2010) by phosphorylating MAPs such APC that control microtubule dynamics. When GSK3 activity is inhibited, APC binds to a microtubule plus end, by which it anchors spindle microtubules to the kinetochore and astral microtubules to the cell cortex (Hur and Zhou, 2010). By regulating microtubule assembly, GSK3 signaling is a key determinant of neuronal polarity and axonal extension. Indeed, inhibition of GSK-3β by small molecule inhibitors, peptide inhibitors, or shRNA induced multiple functional axons (Jiang et al., 2005). We found that GSK-3β accumulated in the axons of P3 DRG neurons from *Lis1*^*+/−*^ mice, with a significantly lower ratio of inactive to total GSK-3β (pS9-GSK-3β/GSK-3β) and a significantly higher ratio of active to total GSK-3β (pY216-GSK-3β/GSK-3β) compared to WT mice, indicative of greater GSK-3β activity (Dudek et al., 1997; Hughes et al., 1993). Retrograde transport of GSK-3β was significantly lower in *Lis1*^*+/−*^ DRG neurons than in *Lis1*^*+/+*^ DRG neurons, whereas anterograde transport was similar, which could account for the abnormal GSK-3β accumulation at the axon tip. We speculate that accumulation of GSK-3β at the axon tip impairs cytoskeletal dynamics, limiting axonal extension.

Selective elimination of axons, axon collaterals, dendrites, dendritic branches, and synapses without loss of the parent neuron occurs during normal development and in response to injury or degenerative diseases in the adult brain. Widespread overproduction or overextension of axonal projections, dendritic branches, and synaptic connections requires both small-scale and large-scale pruning to establish precise connectivity, and these same or similar growth and pruning mechanisms may be reactivated for neural plasticity in the adult nervous system (Luo and O’Leary, 2005). For example, the axons of retinal ganglion cells initially overshoot their future termination zone in the superior colliculus. Later, axon segments distal to the termination zone are pruned through local degeneration (Feldheim and O’Leary, 2010). In developing rats, great numbers of fibers extending through the fornix initially grow well beyond the mammillary bodies and into the mesencephalic and pontine tegmentum. This postmammillary component of the fornix is almost completely eliminated during the first few postnatal weeks (Stanfield et al., 1987). Remaining improper connections or excessive pruning may result in neuropsychiatric diseases, such as schizophrenia, depression, attention deficit/hyperactivity disorder, and autism (Liston et al., 2011; Rosenthal, 2011; Saugstad, 2011). It will be of great interest to explore whether regulators of motor proteins and microtubule organization are involved in neuropsychiatric diseases via effects on neural circuit pruning.

While the intrinsic regenerative potential of axons may be partially suppressed for efficient pruning in the maturing nervous system, this decreased potential limits regeneration following traumatic injury and in pathological conditions such as Alzheimer’s disease, Parkinson’s disease, multiple sclerosis, and amyotrophic lateral sclerosis. A previous report indicated that calpastatin induction facilitated regeneration of damaged neurons. Consistent with this finding, the calpain inhibitor SNJ1945 enhanced sciatic nerve regeneration after injury via augmentation of LIS1. Therefore, our study identifies a novel therapeutic approach to peripheral nerve injury.

## Materials and Methods

### DRG and Cortical Neuron Preparation, Culture, and Imaging

Dorsal root ganglia from P3 and P15 mice were dissociated using a previous method (Lindsay, 1988) with modifications. The cells were plated onto Matrigel-coated dishes (Corning, NY, USA) and cultured in DMEM (Wako Chemicals) supplemented with 10% heat-inactivated fetal bovine serum (Nichirei Biosciences), 10 ng/mL 2.5S mNGF (Sigma-Aldrich, St. Louis, MO, USA), and 5 mM uridine/deoxyfluorouridine (Sigma-Aldrich) for 48 h. Neurons were then transfected with the eGFP-LIS1 expression vector using the Neon Transfection System (Invitrogen). Mouse cortical neurons were isolated from P3, P15, and P60 mice. Briefly, cortical tissue was dissected and dissociated by trypsin digestion and trituration in serum-free medium (Hilgenberg and Smith, 2007) and maintained in Neurobasal medium supplemented with B27 (Invitrogen), GlutaMAX (Invitrogen), and penicillin/streptomycin. Fusion constructs of *GSK-3*β with *td-Tomato* or *eGFP* were transfected into DRG neurons to image GSK-3β migration. Particles in axons were tracked using an IX70 inverted microscope (Olympus) equipped with a stage cell incubator held at 37°C (MATS-LH, Tokai Hit). The images were captured with a digital CCD camera (EM-CCD C9100-13, Hamamatsu Photonics) and analyzed using MetaMorph software (MDS Analytical Technologies).

### Immunoblotting and siRNA

Cells or tissues were lysed in phosphate-buffered saline containing 0.2% NP-40. For immunoblotting, proteins were separated by SDS-PAGE under reducing conditions, followed by electrophoretic transfer to PVDF membranes. Membranes were probed using antibodies against βIII-tubulin (Abcam), CTCF (Abcam), GAPDH (Abcam), and LIS1, followed by visualization using a secondary antibody conjugated to alkaline phosphatase or horseradish peroxidase. Blots were developed using the BCIP/NBT phosphatase substrate system (Roche, Basel, Switzerland) or enhanced chemiluminescence technique (Amersham ECL Prime Western Blotting Detection Reagent RPN2232, GE Healthcare) on a LAS-3000 lumino-image analyzer system (GE Healthcare Biosciences, UK). Deprotected and double-stranded 21-nucleotide RNAs targeting mouse CTCF were synthesized by Sigma-Aldrich (Japan).

### Immunocytochemistry

Cells were fixed with 4% (w/v) paraformaldehyde for 15 min at room temperature and permeabilized using 0.2% Triton X-100 for 5 min at room temperature. The cells were then blocked using 5% (w/v) BSA in PBS and incubated with anti-βIII-tubulin (Abcam),anti-LIS1, anti-GSK-3β (BD Biosciences), anti-pS9-GSK-3β (CST), and anti-pY216-GSK-3β (BD Biosciences), followed by incubation with Alexa 488-conjugated anti-mouse IgG, Alexa 555-conjugated anti-rabbit IgG, and/or Alexa 647-conjugated anti-rabbit IgG (Molecular Probes) as appropriate. Nuclei were counterstained using 100 nM 4′,6-diamidino-2-phenylindole (DAPI). Each incubation was performed for 1 h at room temperature. Slides were mounted in FluorSave Reagent (345789, Calbiochem). Immunofluorescence was measured under a laser scanning confocal microscope (TCS-SP5, Leica, or LSM 700, Carl Zeiss) under the control of accessory software (LAS AF, Leica, or ZEN 2012, Carl Zeiss). Nuclei were labeled with DAPI (Thermo Fisher Scientific).

### Generation of Luciferase Reporter Constructs, Transient DNA Transfection, and Luciferase Reporter Assays

A BAC clone carrying murine *Lis1* was obtained from Advanced Geno Techs (Japan). The *Lis1* minigene was cloned into the luciferase reporter vector *pGL4.23* (Promega). *Luciferase* was conjugated in-frame to the end of *Lis1* exon II. For luciferase reporter assays, approximately 5 × 10^4^ DRG neurons per well were seeded in 6-well plates and cotransfected with luciferase constructs and *Renilla* control reporter vector (phRL-TK, Promega) at a ratio of 10: 1 by electroporation using the Neon Transfection System (Invitrogen). Twenty-four hours after transfection, cells were lysed with Passive Lysis Buffer (Promega), and luciferase activity was measured using the Dual-Luciferase Assay System (Promega).

### Surgical Procedure and Tissue Processing

All mouse experiments were performed with the approval of the Animal Care and Ethics Committee of Osaka City University (authorization number: OCU-08033). First, animals were anesthetized by inhalation of sevoflurane (Wako Chemicals) and intraperitoneal injection of somnopentyl (Kyoritsu). The left SN was exposed from where it emerges over the external obturator muscle from the sciatic notch to the trifurcation above the popliteal fossa. The nerve was sharply transected 5 mm proximal to the sciatic trifurcation, the ends were left to retract *in situ*, and the wound was closed (Alant et al., 2013). One week (6 mice), one month (6 mice), and six months (5 mice) after transection, the incision site was opened and three approximately 1 mm long segments of the nerve were obtained for preparation of transverse sections. The nerve segment was first fixed for a few seconds in a small drop of solution containing 2.5% glutaraldehyde and 0.5% sucrose in 0.1 M Sorensen phosphate buffer (pH 7.4) to stiffen the tissue for correct orientation and then placed in the same fixative solution for 6–8 hours. SN segments were then dehydrated through a graded ethanol series (50%, 70%, 80%, 90%, and 100%) and embedded in Epon 812 resin, followed by thin sectioning at 70 nm thickness using an Ultramicrotome EM UC-6 (Leica Microsystems, Vienna, Austria). The specimens were finally stained with 0.4% lead citrate and observed under a transmission electron microscope (Hitachi).

### Functional Analysis: Walking Tracks

Before SN injury and once weekly for 5 weeks thereafter, all animals were subjected to walking-track analysis based on the protocol described by Inserra et al. (1998). Paw prints were recorded by painting the hind paws with India ink and animals were tracked as they walked along a 45 × 6.5 cm sheet of white paper (Canson A4, 140 g/m2). The paw prints of untreated and SNJ1945-treated mice were analyzed for two parameters: (1) toe spread (TS) as measured by the distance between the first and fifth toes and (2) print length (PL), the distance between the third toe and the hind pad. SFI was calculated according to the formula of Inserra et al. (1998):

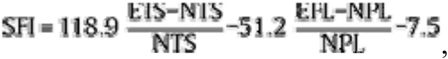

where TS is the toe spread in mm, PL is the print length in mm, and E and N indicate the experimental and normal hind foot, respectively. Differences between the groups were assessed using the Mann–Whitney *U* test. A *P* value < 0.05 was considered statistically significant.

### *In Utero* Electroporation and Histological Examination

*In utero* electroporation was performed using pregnant *Fezf2-Gfp* BAC transgenic mice as described previously (Gong et al., 2003; Kwan et al., 2008) with minor modifications. Approximately 1 μL of the plasmid solution (0.3 μg p*CAG-td-Tomato*, 0.3 μg p*CAG-Lis1*, 0.3 μg p*CMV-CTCF*, or 0.3 μg shRNA against *CTCF* (Sigma-Aldrich) with 0.03% fast green) was injected into the lateral ventricle of intrauterine embryos at embryonic day 12.5 (E12.5). The head of the embryo was placed between the disks of a forceps-type electrode (3 mm disk electrodes, CUY650P3; NEPA GENE, Chiba, Japan) and electronic pulses (30−50 V, 50 ms, five times) were applied for gene transfection into the cerebral wall.

After *in utero* transfection, P15, P18, and P21 mice were perfused by periodate-lysine-paraformaldehyde fixative, pH 7.4. The mouse brains were removed and immersed in the same fixative overnight at 4°C. After fixation, the brains were placed in a 20% sucrose solution, embedded in OCT compound (Sakura), and frozen in liquid nitrogen. The frozen blocks were cut with a cryostat into 16 μm thick sections. Immunofluorescence analyzes were conducted under a laser scanning confocal microscope (TCS-SP5, Leica, or LSM 700, Carl Zeiss) using the accessory software (LAS AF, Leica, or ZEN 2012, Carl Zeiss).

## Acknowledgements

We would like to thank Dr. Kazuhiko Igarashi and Dr. Hiroshi Kiyama for the discussion and suggestions. We also thank Senju Pharmaceutical Co., Ltd., for providing SNJ1945. We are grateful to Yukimi Kira and Yoriko Yabunaka for the technical support and Hiromichi Nishimura and Keiko Fujimoto for mouse breeding. This work was supported by a Grant-in-Aid for Scientific Research from the Ministry of Education, Science, Sports and Culture of Japan to Shinji Hirotsune. This work was also supported by the Uehara Memorial Foundation, the Naito Foundation, and the Takeda Science Foundation to Shinji Hirotsune.

### Competing Interests

We declare no competing interests.

### Authors’ Contributions

K. Kumamoto and S. Hirotsune designed the study, planned and performed most of the experiments, analyzed data, and wrote the manuscript. T. Iguchi and M. Sato contributed to *in utero* experiments. T. Uemura contributed to behavior.

**Figure S1 (related to Figure 1): Whole-neuron images showing LIS1-dependent axonal extension.**

(A) Overview of Tuj1 staining of DRG neurons isolated at various developmental stages as indicated (left). The majority of DRG neurons from WT P3 mice displayed robust axonal extension, whereas axonal extension was clearly limited in WT P10, WT P15, and *Lis1*^+/−^ P3 neurons. (B, C) Overview of Tuj1 staining in DRG neurons isolated from P3 (B) or P15 (C) WT mice overexpressing LIS1 by eGFP-*Lis1* transfection. Exogenous expression of LIS1 augmented axonal extension in DRG neurons from WT P15 mice. (D) Overview of SNJ1945 treatment. Administration of SNJ1945 facilitated axonal extension in DRG neurons from WT P15 mice. (E) LIS1 expression after SNJ1945 treatment. GAPDH was used as the internal control for western blotting. SNJ1945 markedly enhanced LIS1 expression of DRG neurons from P15 but not from P3 mice. Relative intensities from densitometric analysis 24 hours after plating are shown in the right panel. The ratio of LIS1/GAPDH in control P3 mice is defined as 1.0. ∗∗*P*< 0.01 by Student’s *t*-test.

**Figure S2 (related to Figure 3): Whole-neuron images showing CTCF effects on axonal extension.**

(A, B) Overview of Tuj1 staining in WT DRG neurons isolated at P3 (A) or P15 (B). eGFP-*CTCF* transfection suppressed axonal extension in WT P3 DRG neurons, whereas there was a little effect on WT P15 DRG neurons. td-Tomato-*Lis1* cotransfection rescued suppressed axonal extension by eGFP-*CTCF* (A).

**Figure S3 (related to Figure 4): The effect of CTCF depletion by siRNA.**

(A) Overview of Tuj1 staining in WT DRG neurons isolated at P3 or P15. Depletion of CTCF by siRNA facilitated axonal extension in DRG neurons from WT P15 mice.

**Movie S1**

Left: eGFP-tagged GSK-3β movement in P3 DRG neurons from *Lis1*^*+/+*^ mice.

Middle: eGFP-tagged GSK-3β movement in P3 DRG neurons from *Lis1*^*+/−*^ mice.

Right: eGFP-tagged GSK-3β movement in P3 DRG neurons from SNJ1945-treated *Lis1*^*+/−*^ mice.

**Movie S2**

Locomotor behavior of a hemiaxotomy mouse without SNJ1945 treatment.

**Movie S3**

Locomotor behavior of a hemiaxotomy mouse after SNJ1945 treatment. SNJ1945 facilitated recovery of paw grip on the axotomy side.

